# Parallel maturation of hippocampal memory and CA1 task representations

**DOI:** 10.1101/2024.03.21.585945

**Authors:** Juraj Bevandić, Federico Stella, H. Freyja Ólafsdóttir

## Abstract

Hippocampal-dependent memory is known to emerge late in ontogeny and its full development is protracted. Yet, the changes in hippocampal neuronal function that underlie this delayed and gradual maturation remain relatively unexplored. To address this gap, we recorded ensembles of CA1 neurons while charting the development of hippocampal-dependent spatial working memory (WM) in rat pups (∼2-4weeks of age). We found a sharp transition in WM development, with age of inflection varying considerably between individual animals. In parallel with the sudden emergence of WM, hippocampal spatial representations became abruptly task specific, remapping between encoding and retrieval phases of the task. Further, we show how the development of task phase remapping could partly be explained by changes in place field size during this developmental period as well as the onset of precise temporal coordination of CA1 excitatory input. Together, these results suggest that a hallmark of hippocampal memory development may be the emergence of contextually specific CA1 representations driven by the maturation of CA1 micro-circuits.

---

The hippocampus supports the ability to anchor memories for events in our lives to their spatial-temporal context (episodic memory^1^). A capability central to adaptive decision-making and planning for the future. Yet, hippocampal-dependent memory is known to develop late – with the first two years in human life characterized by a near absence of episodic memory^2^ – and its full maturation is protracted^3,4^. In recent years, scientists have begun elucidating the ontogeny of hippocampal neuronal representations. These studies have shown that hippocampal place cells^5^ – the primary cellular model for hippocampal memory – can already be recorded early in third postnatal week^6,7^; before the first signs of hippocampal memory emerge^8-10^. However, although present, various lines of evidence suggest place cell function during this period may still be relatively non-specific^11,12^. Further, the developmental emergence of non-spatial place cell encoding – such as of different trajectories^13,14^ or task phases^15^ – has also not been explored. The ability to integrate diverse cues may be critical to the ability to form detailed and specific memories – a hallmark of hippocampal memory. As such, we sought to investigate the development of non-spatial encoding in hippocampal place cells.

To this end, we recorded from populations of CA1 place cells in P17-P28 rat pups – the developmental period when adult-like hippocampal memory emerges in rats^8-10,16^-while the pups carried out a hippocampal-dependent memory task that requires animals to alternate between sample and memory-guided task phases (discrete trial, delayed non-matching to place (DNMP) ^17,18^). In adult animals, this task is associated with a strong influence of non-spatial features of the task on CA1 place cell activity – place cells display robust remapping between the two phases of the task^15^. Here we sought to chart the ontogeny of such task phase remapping in relation to the maturation of hippocampal memory.

We recorded from 7-70 CA1 place cells per session in parallel (mean = 17.11 (SD=9.02), see Online Methods) from 13 pups while the pups carried out the DNMP task in a T-maze (Figure 1A, Online Methods). Experiments started between P17 and P24 and continued every day for approximately one week. Between P17 and P28 we observed a gradual improvement in the pups’ average performance on the task (r = 0.97, p < 0.0001), reaching eventually performance comparable to adults (*t*(118) = 0.42, p = 0.68). This is in agreement with previous work that showed pups can reliably carry out this task, and other hippocampal-dependent tasks, at around 3weeks of age^8,9,19^. However, previous studies have also highlighted that the ability to carry out tasks similar to the DNMP emerges abruptly in individual animals^20^. To address this question, we fitted sigmoid curves to individual pups’ developmental curves (Figure 1C, Online Methods). We found sigmoid fits captured the developmental curves significantly better than linear fits (AIC sigmoid fits = -37.98 (SD=6.88), linear fits = -34.26 (SD=6.27), *t(12)* = -2.78, p = 0.0.017, Figure S1C), suggesting the developmental emergence of this form of hippocampal memory occurs abruptly (i.e. overnight). Further, the timing of inflection points varied notably between animals (Figure 1C,D), with the earliest inflection point observed at P19 and the latest at P24. Importantly, this abrupt improvement in performance could not simply be explained by experience, as animals that started experiments in the post-weaning period (>P21) already performed above chance at day1 and showed a significantly shallower sigmoid fit (2-sample Kolmogorov-Smirnov test, p = 0.028, Figure 1E, Figure S1A). Further, we did not observe a difference in development curves for pups who were weaned *after* experiments concluded as opposed to at P21 (*t*(11) = 0.18, p = 0.86, Figure S1B).

**Figure 1.**
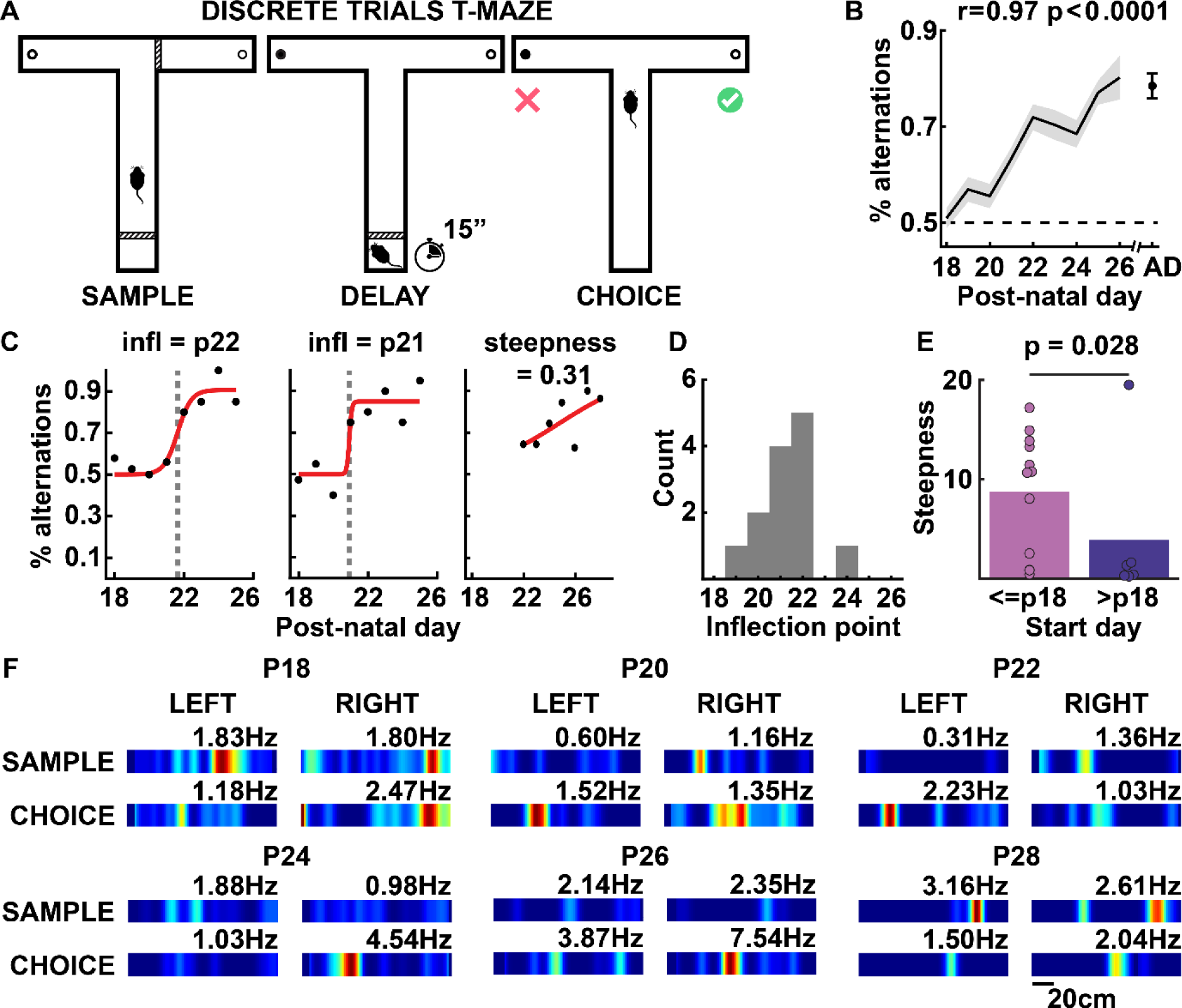
Hippocampal-dependent memory develops abruptly. (A) Schematic of the DNMP task. (B) Mean percentage of correct alternations of pups across age. Shaded area shows SEM. (C) Fitted sigmoids to individual animal developmental curves (black cricles show performance on individual days, red line the fitted sigmoid and the grey dashed line the inflection point). (D) Frequency distribution of age (PD) of inflection. (E) Sigmoid steepness for animals that started the day at P18 and those that started the experiment when older. Circles show individual animal slopes. (F) Example linearized ratemaps from ages divided into the four different run types.

Is this abrupt behavioural development supported by changes in neuronal activity? In the first instance, we assessed how CA1 spatial encoding changed during this developmental period and how, or if at all, it relates to the development of the animal’s ability to carry out the task. To this end, we correlated ratemaps for the two arms of the T-maze (Figure 2A(i), Online Methods). Spatial correlations between ratemaps of the two arms were generally low (mean = 0.07 (SD=0.22), Figure 2A(ii)), suggesting place cells differentiated between the two geometrically identical arms. To ensure the low spatial correlations could not merely reflect unstable place coding, we correlated ratemaps for odd and even runs on the same arms (Online Methods) and compared the distribution of these stability scores against the left vs right arm remapping scores (Figure 2A(ii)). We found the remapping scores were significantly lower than the stability scores (mean = 0.30 (SD=0.15), p < 0.0001, 2-sample Kolmogorov-Smirnov test), indicating the low correlations between the left and right arm ratemaps indeed reflect remapping. We next asked if spatial remapping changed in tandem with the developmental improvements observed for individual animals on the DNMP task. To this end, we correlated the average session spatial remapping scores against days to/from inflection (0 indicates first day after inflection). Spatial correlations scores did not correlate with the development of DNMP memory (r = - 0.09, p = 0.47, Figure 2A(iii)). Indeed, at the earliest ages tested CA1 cells showed reliable remapping between the two arms (r = 0.052 (SD=0.18), *t(5)* = -3.05, p = 0.029, 1-sample t-test P17-P18 remapping vs stability), consistent with previous research^12^, and the remapping observed did not differ from adult spatial remapping (r = 0.18 (SD = 0.20), *t(86)* = -1.44, p = 0.16, Figure 2A(iii)).

**Figure 2.**
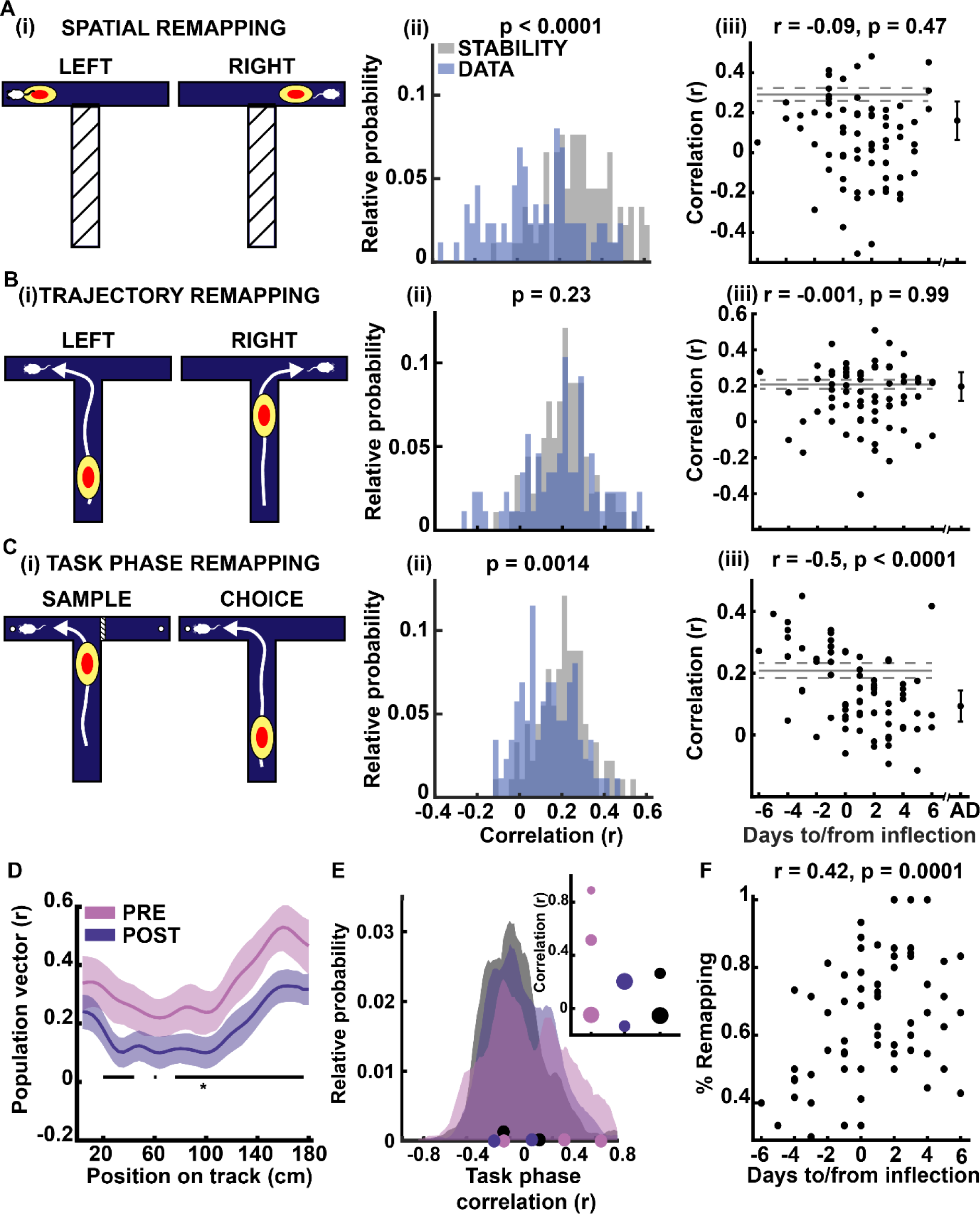
Task phase remapping ontogeny predicts maturation of hippocampal memory. (A) (i) Spatial remapping was computed by correlating place cell ratemaps for left and right arms. (ii) Distribution of average session remapping scores (blue) and stability scores (grey). (iii) Mean session remapping scores as a function of days to/from inflection point. Horizontal grey line shows mean stability score along with standard deviation (dashed line). (B,C) Same as A but for trajectory and task phase remapping, respectively. (D) Population vector correlation between sample and choice ratemaps for pre and post inflection periods. Shaded area shows 95% confidence intervals from bootstrapping data. Black line underneath PV indicates bin where there is a significant difference between pre and post inflection periods. (E) Distribution of task phase remapping scores for pre (light purple) and post (dark purple) inflection periods as well as adults (grey). Circles on the x-axis show the centre of fitted gaussian components. Inset: Centre of gaussian components for pre and post inflection periods and for adults, y-axis shows the task phase ratemap correlation that corresponds to the centre of individual components. The size of the circle shows the proportion of data that is captured by individual components. (F) Proportion of cells remapping between choice and sample trials across inflection points.

As noted previously, studies in adult rodents have shown significant trajectory (left-vs right-bound trials) and task phase (sample vs choice trials) remapping in the DNMP task. Next, we sought to chart their ontogeny. Starting with trajectory remapping, we found no evidence for this form of remapping in our data (Figure 2C(i)); the average correlation between right- and left-bound stem ratemaps did not differ from the stability distribution (p = 0.23, 2-sample Kolmogorov-Smirnov test, Figure 2C(ii)). Further, the degree of trajectory remapping did not show any relationship with the DNMP developmental curves (r = -0.001, p = 0.99, Figure 2C(iii)). On the other hand, we found the CA1 cells remapped strongly between the sample and choice trials (p = 0.0014, Figure 2D(i-ii)), and the degree of task-phase remapping correlated significantly with days to/from inflection point (r = -0.5, p < 0.0001), becoming comparable to adult task phase remapping in the post-inflection period (*t*(58) = 0.15, p = 0.88). To note, we also correlated the different measures of remapping to post-natal age (Figure S2A-C). These correlations were notably weaker than those observed for inflection point (trajectory remapping: r = -0.13, p = 0.25; task phase remapping: r = -0.22, p = 0.04), highlighting the need to account for individual variation when studying the neuronal basis of cognitive development.

Importantly, the relationship between task-phase remapping and DNMP developmental curves could not be explained by experience. A partial correlation controlling for the effect of experience had only a small effect on the inflection point vs task-phase remapping correlation, and it remained robustly significant (r = -0.47, p < 0.0001). Indeed, using a general linear model to predict task-phase remapping from experience, post-natal age and days to/from inflection (Online Methods) showed that only days to/from inflection could significantly predict developmental changes in task-phase remapping (GLM: inflection point: *t*(71) = -4.56, p < 0.0001; post-natal day: *t*(71) = 0.94, p = 0.35; experience: *t*(71) = 1.2, p = 0.24). Finally, we found movement speed did not differ between the two trial phases (choice = 21.24cm/sec (SD=10.87), forced = 19.01cm/sec (SD=10.88), *t*(172) = 1.35, p = 0.18), although median speed increase significantly with age (r = 0.52, p < 0.0001). Importantly, task phase remapping remained a significant predictor of days to/from inflection after controlling for age-related changes in speed (r = -0.38, p = 0.0011).

To corroborate these findings and to explore where remapping occurred on the maze, we turned to a population vector analysis (Online Methods). For each spatial bin (4cm) we correlated the population vectors between sample and choice ratemaps for each session, and then computed the average correlation across all sessions recorded during the pre- and post-inflection period. In agreement with the remapping analysis described above, we found population vector correlations were significantly lower during the post inflection period compared to the pre inflection period (Figure 2D). To note, dividing the post-inflection population vector correlations in two (peri: inflection points 0-2; post: >2 days post inflection) showed no further changes in remapping (all spatial bins p > 0.05, Figure S3). This suggests task-phase remapping emerges abruptly in development, mirroring the abrupt development of hippocampal memory.

Next, we sought to characterize whether developmental changes in remapping reflect homogeneous changes in place cell task phase coding across the CA1 population. To this end, we fitted gaussian distributions to the PRE and POST task phase remapping distributions (Online Methods). During the pre-inflection period, we found that the task phase remapping distribution was best fitted with three Gaussian components (AIC = 432.35), and the three components were centred on r = -0.05 (66%), r = 0.52 (30%) and r = 0.89 (3.6%) correlation scores (Figure 2E). This suggests heterogeneous task phase encoding during the pre-inflection period, with some cells remapping while others did not. During the post-inflection period, however, the task phase correlation distribution could be captured by only two components (AIC = 230.97), one centred on r = -0.13 (30%) and another on r = 0.20 (70%, Figure 2E). Thus, the high correlation component (r = 0.89) observed during the pre-inflection period disappeared post inflection, and only two sub-populations – both centred on low correlation scores - of task phase coding were apparent in the population, suggesting task phase remapping had become nearly ubiquitous. In agreement with this, we found days to/from inflection could be reliably predicted by the proportion of cells that showed task phase remapping (r = 0.42, p = 0.0001, Figure 2F). To note, fitting gaussian distributions to adult task phase remapping distributions (Figure 2E), also revealed two components (AIC = 47.75) centred on similarly low correlation coefficients of r= -0.05 (71%) and r = 0.26 (29%) as during post-inflection. This underscores the observations that task phase coding emerges abruptly in development.

These findings led us to ask what could explain the sudden developmental emergence of task phase remapping? We first explored whether days to/from inflection could be predicted by changes in place cell rate, place field size or place cell sparsity (Online Methods). Days to/from inflection showed no relationship to place cell activity rate (peak rate: sample: r = 0.19, p = 0.1, choice r = 0.09, p = 0.42, Figure S4A) nor to sparsity (% of cells active: sample = r = -0.21, p = 0.06, choice: r = 0.06, p = 0.62, Figure S4B). However, we observed a significant correlation between the size of place cell’s place field and days to/from inflection (sample r = -0.41, p = 0.0002; choice r = -0.23, p = 0.042; Figure S4C), suggesting spatial coding becomes more precise as cognitive development unfolds. Yet, controlling for developmental changes in place field size could not fully account for the correlation we observed between days to/from inflection and task phase remapping (partial correlation r = -0.41, p < 0.001); suggesting the emergence of task phase remapping may only partially be explained by changes in place field size in development.

An alternative hypothesis is that task phase remapping may reflect maturation in the temporal coordination of input to CA1. CA1 receives primarily two glutamatergic inputs – one from layer three of the entorhinal cortex (ECIII) and another from area CA3. These two inputs are thought to reflect distinct hippocampal network states, supporting complementary processes. ECIII input has been purported to support encoding-related processes while CA3 input may rather support memory-guided processes^21,22^. As the DNMP task requires animals to learn to alternate between encoding and memory-guided phases, perhaps the ontogenetic emergence of task-phase remapping, and thereby hippocampal memory maturation, reflects developmental changes in CA1 input alignment to distinct task phases. Namely, with development CA3 input may become more dominant and specific to the choice phase – which requires an execution of memory-guided actions - while ECIII input may be preferentially dominant during encoding-driven sample phases.

To address this question, we analysed CA1 single-unit activity and LFP markers that provide a proxy for CA3/ECIII input balance in CA1. In the first instance, we analysed theta phase preferences of place cell spikes during the sampling and choice phases. As ECIII input arrives earlier in a theta cycle relative to CA3 input^23^, we reasoned that during the post-inflection period, sample phases should be associated with earlier theta phase spiking compared to choice phases but that no difference in phase preference should be observed in the pre-inflection period. Consistent with this, we found place cells tended to fire near the trough of locally recorded theta-band oscillations during the pre-inflection period (sample mean angle = 131.5°, choice mean angle = 110.4°, Figure 3A(i)), and the phase preference between sample and choice phases of the task did not differ (95% bootstrap confidence intervals = [-0.85, 0.10]). During the post-inflection period, however, we found phase preferences between the two task phases differed reliably (sample = 111.4°, choice = 154.5°, Figure 3A(ii)), with place cells firing at significantly earlier phases of a theta cycle during sample phases of the task relative to choice task phases (95% bootstrap confidence interval = [0.01, 1.42]). Additionally, theta phase locking – thought to derive from CA3 input – decreased significantly as the animals started to be able to perform the task reliably (r = - 0.67, p < 0.001, Figure 3B). Importantly, this effect was driven by the sampling trial phases (p = 0.006, 2-sample Kolmogorov-Smirnov test, Figure 3C). Phase locking during choice trial phases did not change significantly between pre- and post-inflection periods (p = 0.82, 2-sample Kolmogorov-Smirnov test, Figure 3D).

**Figure 3.**
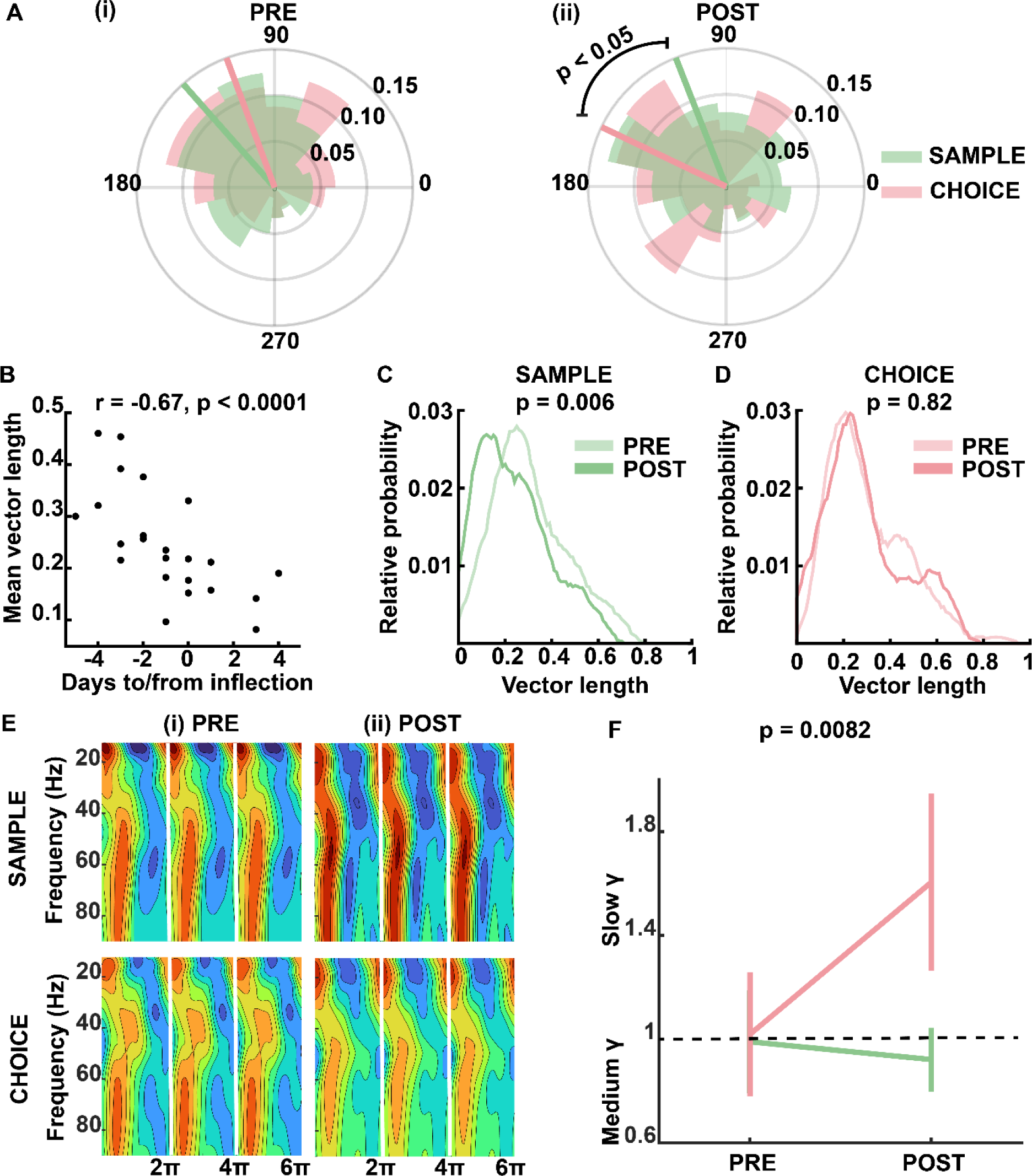
Hippocampal memory maturation is associated with task phase-based coordination of CA1 input. (A) (i) Circular histogram of CA1 cell phase preferences for sample and choice trials during the pre inflection period. Coloured diagonal lines show the circular mean for each trial type. (ii) Same as (i) but for the post inflection period. (B) Mean session phase locking (Raleigh vector length) as a function of days to/from inflection. (C) Frequency distribution of phase locking scores for the pre and post inflection periods for sample trials. (D) Same as (C) but for choice trials. (E) (i) Cross frequency coupling to theta-band oscillations during sample (top) and choice (bottom) trials, pre inflection, x-axis shows phase in a theta cycle. Note, three theta cycles are shown for clarity. (ii) same as (i) but for the post inflection period. (F) Average sample and choice slow/medium gamma coupling ratio for pre and post inflection periods, error bars show 1SD.

Finally, to corroborate these findings we investigated coupling of slow and medium gamma oscillations to CA1-recorded theta-band oscillations in sampling and choice trials and assessed how theta-gamma coupling changed between the pre- and post-inflection periods. Slow gamma (∼20-40Hz), recorded in CA1, is known to derive from upstream CA3 activity while medium gamma oscillations (∼50-70Hz) are thought to originate from ECIII^24^. Thus, measuring the relative coupling of the two gamma-band oscillations to CA1 theta-band oscillations is another approach to assess the relative influence of ECIII and CA3 input over CA1 activity^24,25^. During the pre-inflection period, we found the ratio between slow and medium theta coupling to be comparable during sample and choice phases of the task (sample slow/medium gamma ratio = 0.99 (SD=0.20), choice = 10.2 (SD=1.02), p = 0.48 based on bootstrapped ratio scores, Figure 3E,F(i)). During the post-inflection period, however, we found a robust difference in the relative theta coupling of the two gamma bands between the two task phases (sample = 0.92 (SD=0.12), choice = 1.61 (SD=0.34), p = 0.0066 based on bootstrapped ratio scores, Figure 3E,F(ii)), with the ratio shifted towards medium gamma during sample phases but towards slow gamma during choice phases (Figure 3F). To note, despite changes in movement speed between the two developmental epochs, these changes could not explain the shift in slow-to-medium gamma balance between the sampling and choice trials post inflection (Figure S5, Online Methods). In sum, consistent with the theta phase locking and phase preference analyses described above, it seems that the two excitatory inputs to CA1 become better aligned to the distinct task phases as the animals’ hippocampal memory develops.

Here, we show that hippocampal-dependent memory and CA1 task phase remapping emerge in parallel in ontogeny. As the ability to carry out a spatial working memory matured so did CA1 neuronal representations for specific task phases. These findings suggest that the development of hippocampal memory may be underpinned by the development of functionally specific CA1 neural representations. Contemporary theories of hippocampal memory development propose that one of the hallmarks of memory maturation is that memory becomes less generic and more specific^26^. This study provides, for the first time, insight into the changes in neuronal coding that may underlie this critical cognitive developmental milestone.

Although the origin of this neuro-developmental milestone remains to be ascertained, we propose it may reflect the combined emergence of precise CA1 place fields and adult-like temporal organization of CA1 glutamatergic input. On the one hand, as place fields become smaller, the ability to distinguish between related representations may increase. Alternatively, if with development the two phases of the task become associated with distinct excitatory inputs, this would naturally lead to different fields emerging for the distinct phases. A more tempting hypothesis is that these phenomena are in fact inter-dependent: precise, information-rich, spatial representations might depend on effective integration of separate input streams in CA1, potentially relying on their temporal organization. The structured interaction of sensory-based and memory-based information in CA1 could be fostered by a shift from competing ECIII and CA3 inputs early in life to their spatio-temporal segregation in adults. In turn, such arrangement could provide the framework for the emergence of high-dimensional representations, supporting adult-like memory capabilities. Finally, it remains to be seen what role the medial prefrontal cortex (mPFC) plays in relation to this process. The mPFC, and particularly mPFC- hippocampal interactions, are known to be crucial for accurate DNMP performance in adults and the mPFC develops significantly during this period^27^. Perhaps the ontogeny of specific CA1 representations, and mature circuit function, reflects the maturation of hippocampal-mPFC pathways.

## Methods

### Experimental model and subject details

Thirteen Lister Hooded rat pups (30-52g at implantation) underwent a surgical procedure to implant a microdrive carrying eight or sixteen tetrodes of twisted 17 μm HM-L coated 90% platinum, 10% iridium wire (California Fine Wire), targeting the right CA1 (ML: 2.0-2.3mm, AP: 3.0mm posterior to bregma). Electrode tips were gold plated to reduce impedance to 150-300kOhm at 1kHz. Pups were allowed to recover from surgery housed together with littermates (and dam if pre-weaning) for 48h. The animals always had *ad libitum* access to water and food, and were housed on a reversed 12-h light-dark cycle. Eleven adult Lister Hooded rats (330-400g at implantation) underwent the same procedure as pups. Adult procedures differed from pup procedures in CA1 target coordinates (ML: 2.0-2.3mm, AP: 3.8mm posterior to bregma), one week of post-operative recovery, and they were housed individually.

### Electrophysiological recording

After the post-operative recovery period, electrophysiological activity was screened two to three times a day. All recordings were performed using an Axona recording system (Axona Ltd., St. Albans, UK) which recorded spike-threshold triggered single unit activity, continuous LFP, and position data. Each channel was amplified 5000 – 15000 times and recorded referenced to another channel on a separate tetrode. Spikes and LFP were sampled at 48KHz. Animal position was determined using an overhead infrared (IR) camera recording the location of an array of IR light-emitting diodes (LED) mounted on the headstage. Tetrodes were gradually advanced ventrally in 62.5 – 125μm steps, at least 3h apart, until place cells and sharp-wave ripples were detected.

### Experimental apparatus and procedures

All experiments were performed in a dark room with no natural sources of light. The experimental area of the maze was surrounded by thick opaque black curtains on all sides. The experimental arena consists of a dark brown wooden digit-8 maze (140x140cm) with textured wood running surface. A T-shaped portion of the maze was used for the DTT task, the remaining parts of the maze were blocked off using black metal barriers. A black plastic food well holding 0.1ml of liquid was placed at the end of each arm of the T-maze. In between trials the animals were placed in an inter-trial-interval (ITI) box (12x12cm) at the start of the stem of the Tmaze. Access to the stem was blocked off with a tall removeable barrier. Sleep recordings were conducted in a tall round opaque black plastic box (25x48cm), filled with bedding sand, that was placed on top of the centre of the stem of the maze, directly underneath the IR camera.

Two days prior to experiments, animals were habituated for a single 10–15 minute session per day. Habituation was carried out on the tracks of the digit-8 maze that were unused in the T-maze, such that the physical characteristics of the tracks are identical to the T-maze tracks, but no part of tracks overlapped between habituation and T-maze. During habituation, animals were allowed to self-initiate walking between two ends of a linear track with food wells filled with soy milk at each end. The experimenter refilled the food wells with milk as necessary.

The task was performed in two identical sessions per day, at least 3 hours apart during which time the animals rested in the homecage. The animals were not food nor water deprived, and the conditioning reward was 0.1ml of soy milk formula. Each trial-pair consisted of two runs – SAMPLE and CHOICE. Each SAMPLE trial had access to one of the arms blocked off with a removable barrier, and food was placed in the open arm. The selection of left/right open arms during SAMPLE trials was pseudo-randomised. When the animal reached the end of the open arm and drank its food reward it placed I the ITI box for 15 seconds. The CHOICE trial started after the ITI door was lifted for the animal to walk out into the stem. In this run, both arms were open. The pup got rewarded if they chose the arm opposite to the one rewarded during the SAMPLE trial. After each trial-pair, the animal was placed into the holding box outside the maze for 30-45s intra-trial interval. Sessions lasted between 15-45 minutes. Immediately after finishing each session, provided sufficient single-unit yield, the animals were placed into the sleep box and allowed to rest for an hour.

### Inflection point analysis

To determine developmental inflection points on the DNMP task for individual animals, we fitted sigmoid curves to individual animals’ daily performance data. Specifically, we fit a sigmoid curve to the data using the standard logistic function (Equation 1) where *d* denotes the chance level performance which is set to 0.5, *a* represents the animal’s highest daily performance mean, *c* is the steepness of the curve ranging from 0 to 20, and *b* the inflection point of the sigmoid constrained between the animal’s first and last post-natal day. We used the nonlinear least squares fit option in Matlab R2019b (Mathworks, MA) in order to fit the model to the data. Any animals with fewer than 5 daily means (n = 3 animals) were excluded from the inflection point analysis to ensure a reliable fit.

In cases where the inflection point given by the fitted model falls between two whole numbers, we always rounded up the value. Therefore, the next PD after the inflection point is determined to be day 0. For PRE vs POST analyses, we divided the data into sessions prior to the inflection point (PRE), and sessions from day 0 onwards (POST). For the analysis reported in Figure S4 where we further divided the POST period, PERI was defined as the data from day 0 to day 2, and POST as data form day 3 onwards.

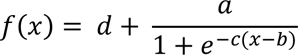

To compare linear and sigmoid fits we computed the Akaike information criterion for linear and sigmoid fits and compared them using a paired samples t-test.

### Place cell analysis

All analyses were restricted to putative principal cells, identified by manual inspection of waveforms across the entire recording session. KlustKwik was applied to spike-thresholded data to sort the data into clusters and then the clusters were manually curated in Tint (Axona Ltd.). We classified spike sorted neurons as place cells by computing Skaggs Information (bits per second), and compared the value we obtained against a null distribution generated by random permutations of spike times. Cells that exceeded 95^th^ percentile of their own shuffle were deemed place cells. Sessions containing fewer than five cells with significant Skaggs information values were excluded from analysis.

To generate ratemaps, spike data was divided into the four different trial types (e.g. SAMPLE left, CHOICE left, etc). Spikes that occurred during stationary periods (<3cm/s) or when the animal was located near the start of the stem (<10cm) were excluded. Next, we linearised animals’ paths, binned dwell time and total number of spikes in 2cm bins, computed firing rates by dividing the binned spikes over binned dwell time, and smoothed them using a Gaussian kernel (sigma= 3 bins).

### Remapping

To analyse remapping between the different trial types we correlated the spatial ratemaps for pairs of runs using the Pearson correlation coefficient (empty bins in both ratemaps were removed). Spatial remapping was performed only on the bins corresponding to the arms of the T-maze. The rest of remapping analyses excluded the arms and were only performed on the bins corresponding to the central stem of the T-maze, which is the only section of the track common to all four run types. Correlations between left- and right-bound ratemaps were refer to as trajectory remapping, while correlations between SAMPLE and CHOICE ratemaps we term task phase remapping. To ensure remapping scores did not just reflect unstable spatial firing, we compared the remapping scores to correlations scores obtained by correlating the ratemaps for odd and even runs for a given trial type. To assess if a particular type of remapping was present in the population we compared the distribution of mean session remapping scores to the stability scores using a 2-sample Kolmogorov-Smirnov test.

To assess if remapping changed with development we correlated average remapping scores obtained in a session with post-natal age/inflection point using a Pearson correlation coefficient. To rule out the effect of experience and changes in median speed during development, we used a partial correlation where the relationship between inflection point and remapping was computed while controlling for the effect of experience/median speed. To compare the relative influence of inflection point, post-natal age and experience we used a General Linear Model (GLM) with remapping scores as the response variable and inflection point, post-natal age and experience as the predictors.

To assess if the proportion of cells remapping between task phases in a session correlated with inflection point we calculated the proportion of cells with task phase remapping scores above 0.25 and correlation these session proportions with inflection point.

### Population Vector Correlation Analysis

To examine the distribution of task phase remapping remapping across the track, we correlated the population vectors for choice and sample trials. Specifically, for each spatial bin (4cm) we computed the Pearson correlation coefficient for z-scored activity of all cells active in a session. The first 10cm at the start of the stem were removed to exclude areas of the maze associated with immobility. The average population vector was then computed for PRE and POST inflection periods and 95% confidence intervals (CIs) computed to assess which spatial bins differed significantly between the two periods. The average PRE and POST population vector correlation was then smoothed with a guassian kernel (sigma = 8cm). To compute the CIs we bootstrapped the session population vector correlations 10,000 times, repeating the analysis separately for PRE and POST inflection periods, for each iteration of the bootstrap we computed the mean population vector correlation. From the bootstrapped data we obtained the 2.5^th^ and 97.th percentile for each inflection period, if the CIs for the two periods did not overlap we deemed the comparison significant. The same procedure was used to compared the population vector correlations between the period immediately after inflection (inflection points 0-2, PERI) and the subsequent days.

### Fitting gaussian components

To fit gaussian components to the distribution of task phase remapping scores during the PRE and POST inflection periods we used the Matlab function fitgmdist, we fitted 1 and 4 components and used the Akaike Information Criterion (AIC) to determine the model with the best fit.

### Place field analyses

Field size and peak rate were assessed by first using the regionprops function in Matlab on rate thresholded ratemaps (bins > 50% of the peak rate of the ratemap). Small fields detected with this method (<20cm long) were removed. Then the area of the field was used as a measure of field size and the highest rate within the field a measure of peak rate. If a cell had multiple fields the average size and peak was computed. To measure sparsity we computed the proportion of all cells recorded there were had a significant skaggs information value in all four run types.

To assess how these place cell features changed with development we correlated them against post-natal age/inflection point using the Pearson correlation coefficient. To control for the effect of field size we used a partial correlation, where inflection point and task phase remapping were the predictor and response variables, respectively, and field size the covariate.

### Theta phase analyses

To analyse theta phase preference and phase locking to theta-band oscillations during SAMPLE and CHOICE trials we first identified the electrode in the CA1 region with the highest power in the theta band (5-12Hz) using LFP data downsampled to 1.2kHz. We performed a wavelet transform to extract the instantaneous phase of the channel’s signal in the theta band. We then used the extracted phase to identify the theta phase of each spike. We filtered out low theta periods where the power of theta was below the mean. Further, we excluded stationary period (<3cm/sec) and only analysed the phase for cells who still had at least 10 spikes within their place field for given run after this filtering.

To compute the phase preference of each cells during SAMPLE and CHOICE runs we calculated the circular mean for each trial type. To assess phase locking we computed the Resultant Vector Length for each cell’s phases. To compare phase preference between SAMPLE and CHOICE runs for the two inflection periods we first boostrapped the preferred phase distributions for each run type 10,000 times. For each iteration of the bootstrap we computed the circular mean of the SAMPLE and CHOICE data. Subsequently, we computed the circular distance between SAMPLE and CHOICE bootstrapped mean phase distributions, and computed confidence intervals based on the returned circular distance distributions. If the CI did not include 0 we concluded that the different between the SAMPLE and CHOICE phases different significantly. To note, analyses were done separately for PRE and POST inflection periods. For phase locking data, we first assessed if phase locking changed with inflection point. To this end, we used a Pearson correlation coefficient between the session mean phase locking scores (for all trial types) and inflection points. We then divided the phase locking data by run type (SAMPLE and CHOICE) and compared the distribution of phase locking scores for each trial type during PRE and POST inflection periods, using a 2-sample Kolmogorov-Smirnov test.

### Theta-Gamma Coupling

To measure coupling of slow and medium gamma oscillations to theta-band oscillations during SAMPLE and CHOICE task phases, we filtered the LFP data so to include only samples where theta power was above the mean and divided the data into SAMPLE and CHOICE periods. We computed the phase-amplitude coupling between theta phase and amplitude of oscillatory components between 15 and 200 Hz: we first extracted the phase of the oscillatory component in the theta band (5-12 Hz). Then we binned theta cycles into 26 phase bins and for each phase bin we computed the average power of the oscillatory components at higher frequencies (>15Hz) obtained from a wavelet decomposition of the full signal. Thus, we obtained a phase-amplitude 2D-matrix of coupling strengths where each value correspond to a particular combination of theta phase interval and degree of amplitude modulation of a faster oscillation. These couplings were then z-scored across each frequency band and average coupling computed for PRE and POST inflection periods and SAMPLE and CHOICE phases. To assess the ratio between slow and medium gamma coupling to theta, we identified the highest coupling observed in the phase-amplitude analysis in the two gamma bands (slow gamma: 18-30Hz, medium gamma: 40-70Hz), and divided the highest coupling observed in the slow gamma band by the highest coupling observed in medium band. Here a value above 1 indicates strongest coupling to slow gamma compared to medium gamma. To assess if slow-to-medium gamma coupling to theta-band oscillations differed between SAMPLE and CHOICE trials during the two inflection periods, we bootstrapped the session cross-frequency coupling spectrograms 10,000 times, and for each iteration of the bootstrap we computed the difference in slow-to-medium gamma coupling. We then computed 95% confidence intervals for the difference score distribution.

To control for the effect of speed during different inflection periods, we repeated the analysis above for different speed bands (low: 3-20cm/s, mid: 20-40cm/s, high: >40cm/s), we then assessed if the difference in slow-to-medium gamma ratios during each inflection period for individual speed bands differed significantly from the original data. To this end, we computed the difference between bootstrapped slow-to-medium gamma ratios during individual inflection periods as described above.

---

**Figure S1.**
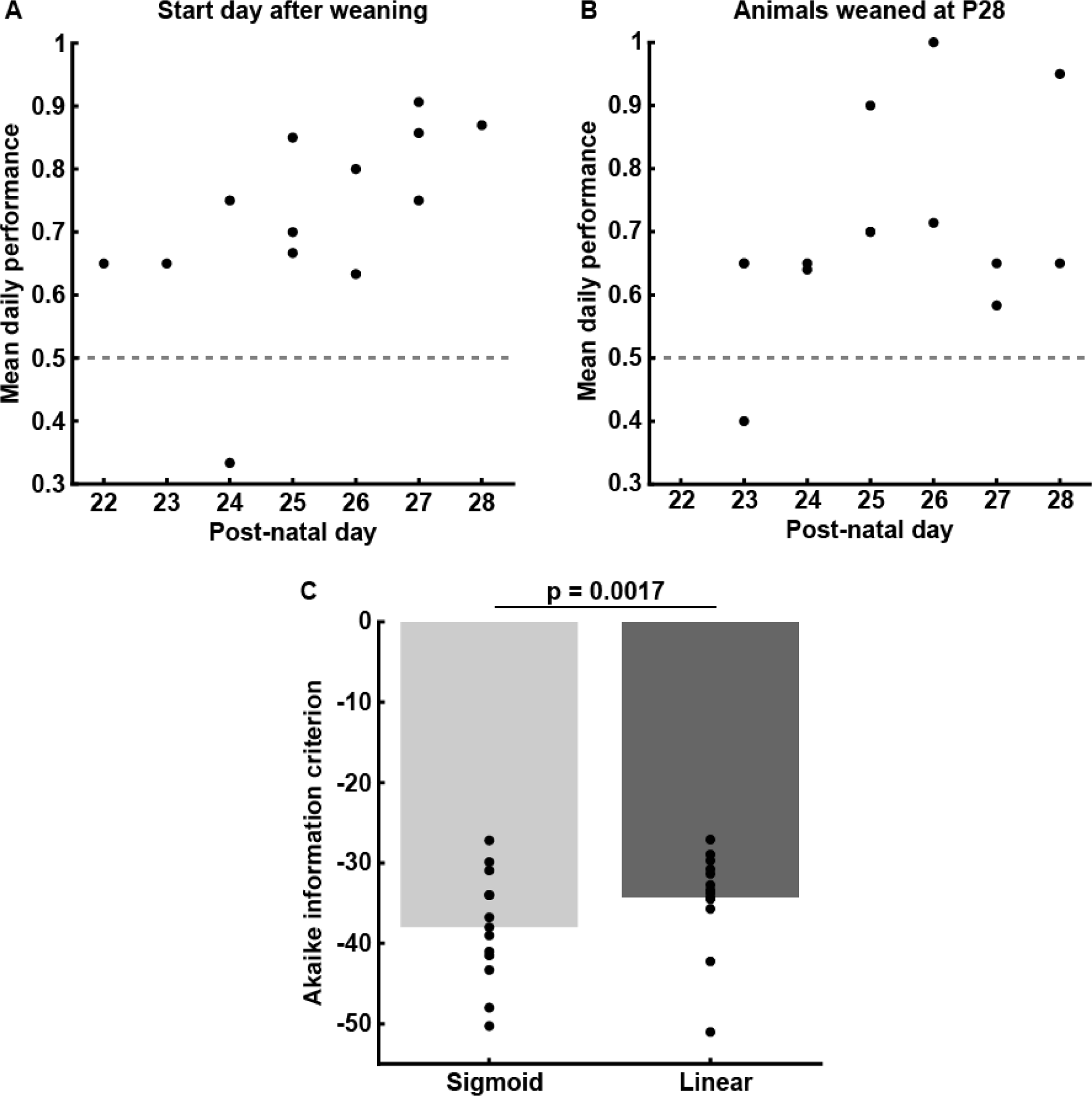
Spatial working memory develops abruptly and maturation is not affected by experience of day of weaning. (**A**) Daily performance of animals that started experiments after weaning (P21). (**B**) Daily performance of pups that were weaned after experiments concluded. (**C**) AIK for Sigmoid (left) and linear (right) fits for individual DNMP developmental curves.

**Figure S2.**
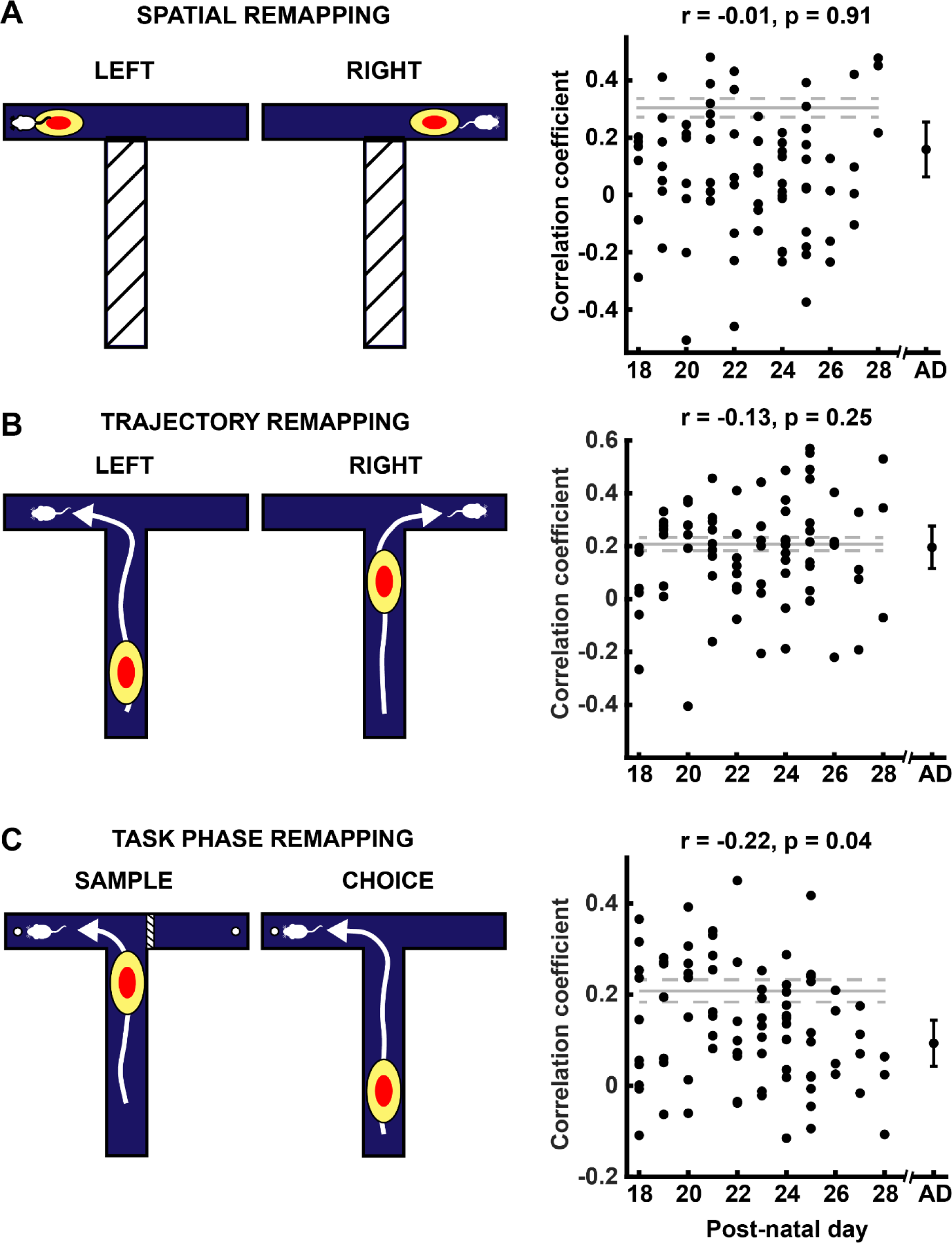
Development of place cell remapping with age. (A) Left: Schematic of spatial remapping analysis (ratemap correlations between left and right arms). Right: Session mean spatial remapping vs post-natal age. (B-C) Same as A but for trajectory (B) and task phase remapping (C).

**Figure S3.**
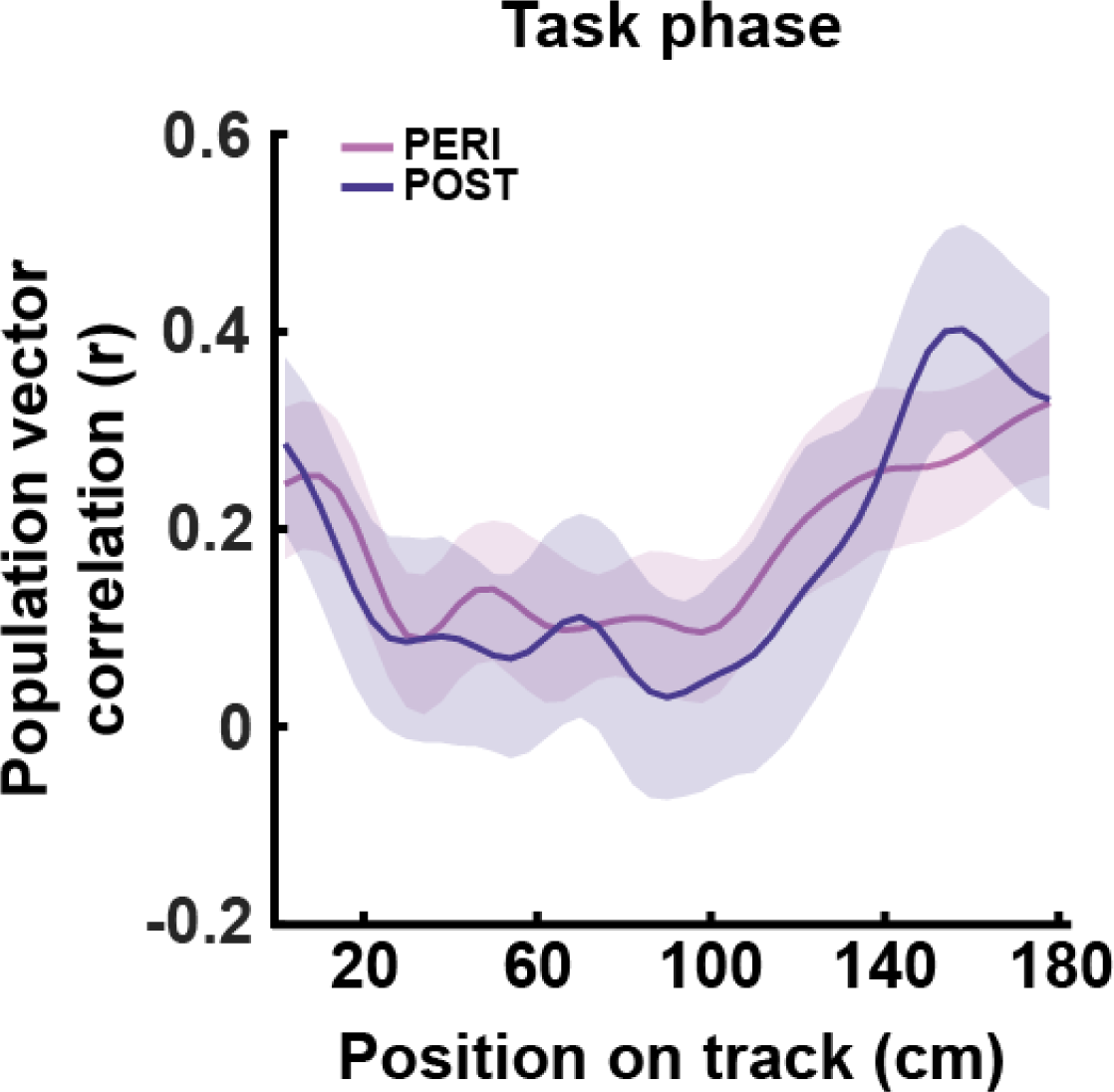
Task phase remapping emerges abruptly following inflection. Population vector correlation between SAMPLE and CHOICE ratemaps for two developmental periods post inflection. PERI=data recorded on the day of the inflection point and two subsequent days, POST = data recorded three or more days after the inflection point. Error bars show 95% confidence intervals.

**Figure 4.**
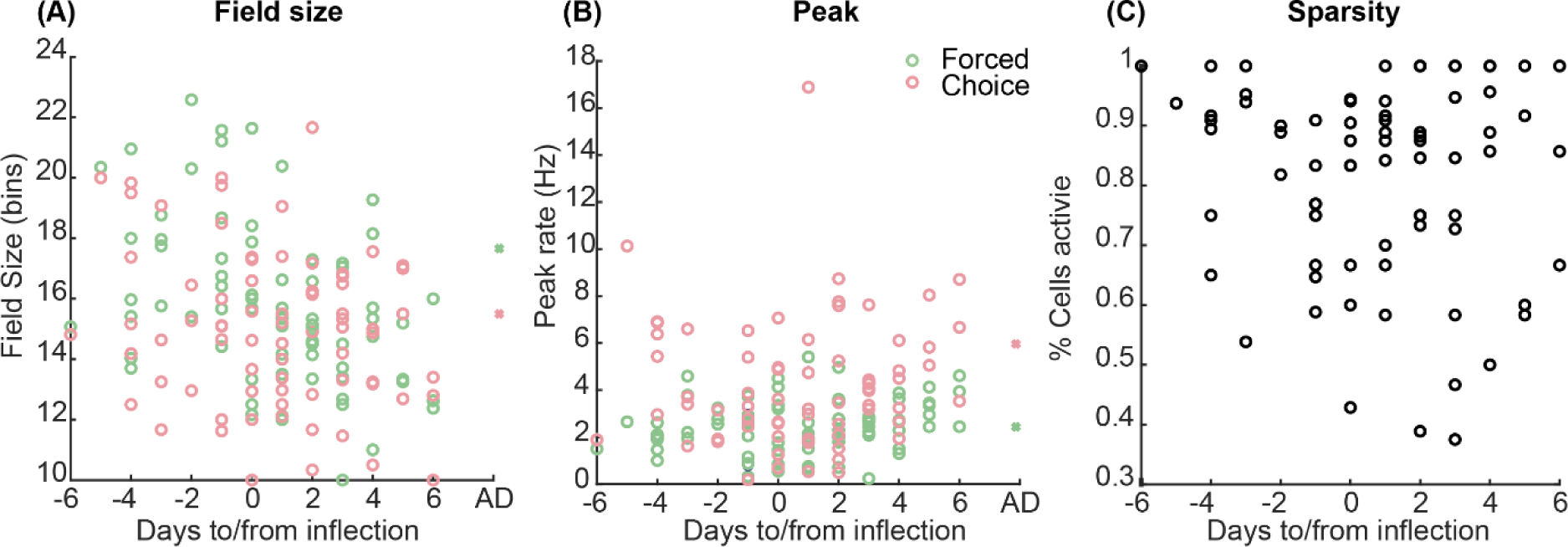
Development changes in place cell activity. (A) Place field size plotted against days to/from inflection. (B-C). Same as A but for peak firing rate (B) and sparsity (C).

**Figure 5.**
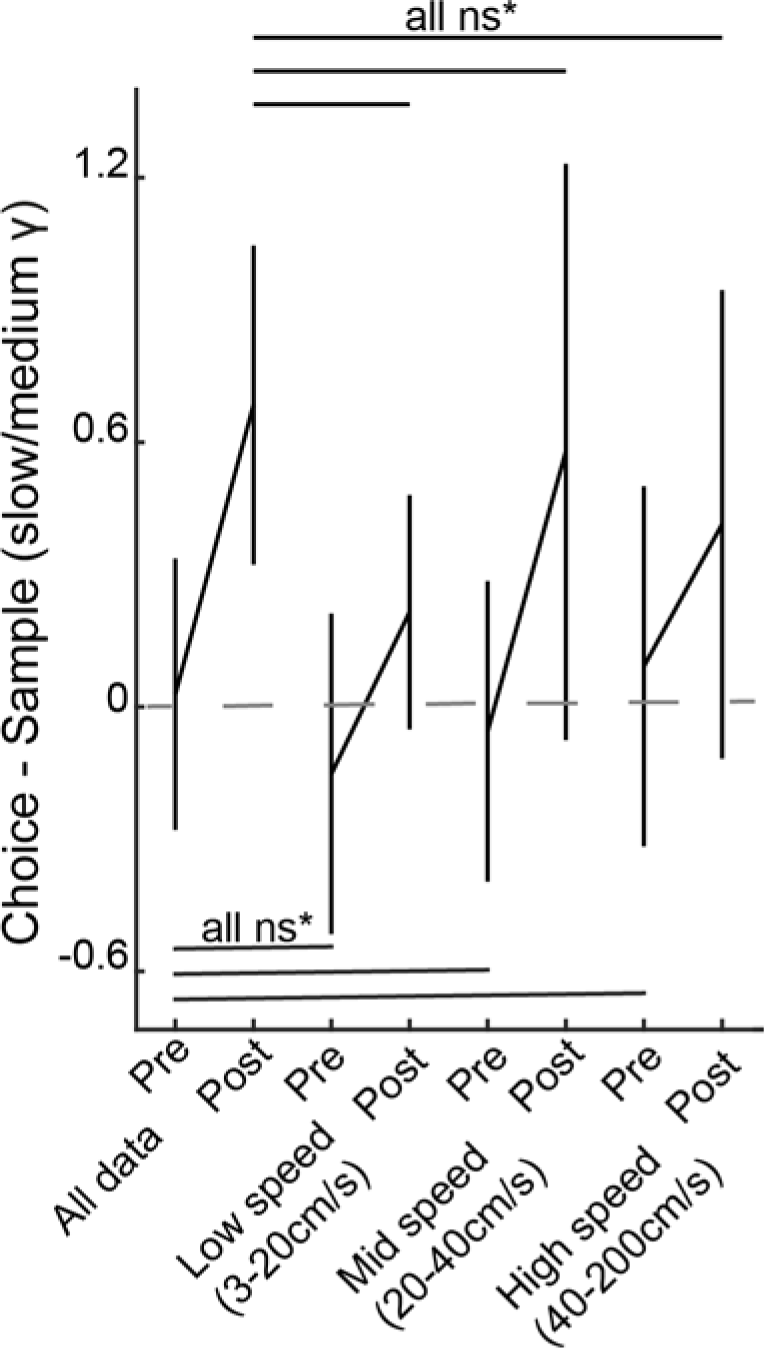
Developmental changes in slow-to-medium gamma coupling to theta is not confounded by speed. Bootstrapped difference scores between slow-to-medium gamma ratios for sample and choice task phases. Far left: original data. Left middle: data filtered for low speed. Right middle: data filtered for medium speed. Far right: data filtered for high speed. The difference scores during pre and post inflection periods for individual speed bands were compared with the difference scores seen in the original data. None of the speed bands revealed difference scores that deviated statistically from the original data.

